# STORIES: learning cell fate landscapes from spatial transcriptomics

**DOI:** 10.1101/2024.07.26.605241

**Authors:** Geert-Jan Huizing, Jules Samaran, Daniele Capocefalo, Anna Audit, Gabriel Peyré, Laura Cantini

**Affiliations:** Institut Pasteur, Université Paris Cité, CNRS UMR 3738, Machine Learning for Integrative Genomics Group, F-75015, Paris, France; CNRS and DMA de l’Ecole Normale Supérieure, CNRS, Ecole Normale Supérieure, Université PSL, 75005, Paris, France

## Abstract

In dynamic biological processes such as development, spatial transcriptomics is revolutionizing the study of the mechanisms underlying spatial organization within tissues. Inferring cell fate trajectories from spatial transcriptomics profiled at several time points has thus emerged as a critical goal, requiring novel computational methods. Wasserstein gradient flow learning is a promising framework for analyzing sequencing data across time, built around a neural network representing the differentiation potential. However, existing gradient flow learning methods cannot analyze spatially resolved transcriptomic data.

Here, we propose STORIES, a method that employs an extension of Optimal Transport to learn a spatially informed potential. We benchmark our approach using three large Stereo-seq spatiotemporal atlases and demonstrate superior spatial coherence compared to existing approaches. Finally, we provide an in-depth analysis of axolotl neural regeneration and mouse gliogenesis, recovering gene trends for known markers as *Nptx1* in neuron regeneration and *Aldh1l1* in gliogenesis and additional putative drivers.

## Main text

Spatial transcriptomics technologies are revolutionizing the study of how cells organize within tissues^1^. Techniques based on high-throughput sequencing have enabled the unbiased discovery of gene expression patterns within their spatial context. For instance, recent studies have revealed previously unknown spatial organization at the tumor-microenvironment interface in melanoma and Alzheimer’s disease amyloid plaque microenvironment^2,3^. The most widely used spatially-resolved sequencing techniques (e.g. 10X Visium) measure spots larger than the typical cell size. However, recent technological developments based on barcoded arrays like Stereo-seq and HDST have reached single-cell resolution, effectively bridging functional and structural characterizations of the cell^4,5^. Recent works have leveraged Stereo-seq to produce large spatiotemporal atlases of various biological processes by profiling a system with spatial transcriptomics at several points in time^4,6,7^. These datasets are ideal for studying cellular dynamics within the tissue during processes such as development and the onset of complex diseases, where cells undergo coordinated transcriptomic changes and spatial reorganization.

Inferring the dynamics of biological processes from single-cell sequencing data requires tailored computational approaches known as trajectory inference methods^8^. Monocle initiated the field of trajectory inference by ordering cells along a pseudotime axis based on their transcriptomic similarities and analyzing gene expression trends along pseudotime^9^. While pseudotime represents the progression along a differentiation process, pseudotime-based methods do not provide a model for the underlying transcriptomic changes, and thus cannot predict a cell’s future transcriptomic state^10^. RNA velocity has thus been proposed to predict changes in gene expression based on splicing dynamics^11^. However, velocity-based methods rely on simple kinetic models that can misinterpret cell dynamics, for instance in the case of transient boosts in transcription^12^.

Multiple methods based on Optimal Transport (OT) have been developed for cases when several time points are available along differentiation. Waddington OT infers trajectories by computing probabilistic cell-cell transitions between adjacent time points^13^. However, it delivers neither a notion of pseudotime nor a notion of velocity. Another class of OT-based methods proposes a continuous model of population dynamics by training neural networks representing a generalized notion of velocity^14^. However, these methods do not order cells along a pseudotime axis. A promising OT-based framework for trajectory inference consists of learning a potential function governing a causal model of differentiation^15–17^. Framing cellular differentiation as the minimization of a potential function is rooted in systems biology and formalizes Waddington’s idea of epigenetic landscape^18,19^. Furthermore, the potential function is a natural alternative to pseudotime, and its gradient yields a rigorous notion of velocity.

The recent development of spatial transcriptomics technologies made it possible to study how space impacts cell trajectories. The addition of spatial information to classical trajectory inference methods is extremely difficult as obtaining a representation for spatial coordinates which is coherent and comparable across all timepoints is challenging. Indeed, spatial coordinates can’t be used as input of the methods the same way gene expression is used because of possible rotations and translations occurring on measured slices. Furthermore, when studying trajectories during development, living organisms can undergo important morphological transformations. A few OT-based approaches inferring trajectories from spatial transcriptomics through time have recently been developed^20–22^. For instance, stVCR learns a spatial velocity along with a gene expression velocity after rigidly aligning the slices. However, this alignment can only partially capture the morphological modifications described above. Additionally both velocity terms in stVCR depend on the time which can allow the model to learn internally a better spatial representation for each time point. This strategy might impair stVCR’s generalization performance on unseen slices of later timepoints. Moreover, SpaTrack learns velocities based on linear OT taking into account both space and gene expression. Moscot computes cell-cell transitions between adjacent time points using an extension of OT called Fused Gromov-Wasserstein^23^ (FGW). FGW is properly suited to compare slices across timepoints since its transport plan is invariant to rotating, translating and rescaling of the spatial coordinates. However, SpaTrack and Moscot only connect cells from adjacent time points, but cannot predict the evolution of cells at unseen future time points.

Here, we propose STORIES, a trajectory inference method capable of learning a causal model of cellular differentiation from spatial transcriptomics through time using FGW. STORIES uses FGW as a machine learning loss to learn a continuous model of differentiation. FGW allows STORIES to implicitly guide the learned potential to depend on the space information without having to use a representation of the space as a direct input. We can therefore learn a differentiation potential shared across all time points and model a general dynamic less prone to overfitting. This potential can then be used to predict the evolution of cells at future time points.

We benchmarked our approach on three large-scale spatiotemporal Stereo-seq atlases, covering mouse development^4^, zebrafish development^6^, and axolotl regeneration^7^ showing superior performances over the state-of-the-art. Furthermore, we used STORIES for the in-depth analysis of cellular trajectories in axolotl neural regeneration and mouse gliogenesis. In both cases, we show that the spatial environment impacts cell fate decisions. We then recover gene trends for known markers, such as *Nptx1* in Nptx+ excitatory neuron regeneration and *Aldh1l1* in gliogenesis. In addition, STORIES uncovers other possible driver genes and transcriptional regulators of cellular differentiation in these contexts, which may be of interest for further biological investigation.

Finally, we provide STORIES as an open-source and user-friendly Python package (github.com/cantinilab/stories). It is based on the Scverse ecosystem, making it easy to interface STORIES with existing tools for single-cell analysis such as Scanpy and CellRank^24–26^. In addition, STORIES benefits from the JAX ecosystem for deep learning and OT computation, enabling the fast handling of large datasets^27,28^.

## Results

### STORIES: a spatiotemporal cell trajectory inference method

We developed SpatioTemporal Omics eneRgIES (STORIES), a tool for single-cell trajectory inference using omics data profiled through spatial and temporal dimensions (github.com/cantinilab/stories). STORIES allows studying dynamic biological processes in their spatial context by identifying cell fates, gene trends, and candidate transcriptional regulators (see Figure 1).

**Figure 1.**
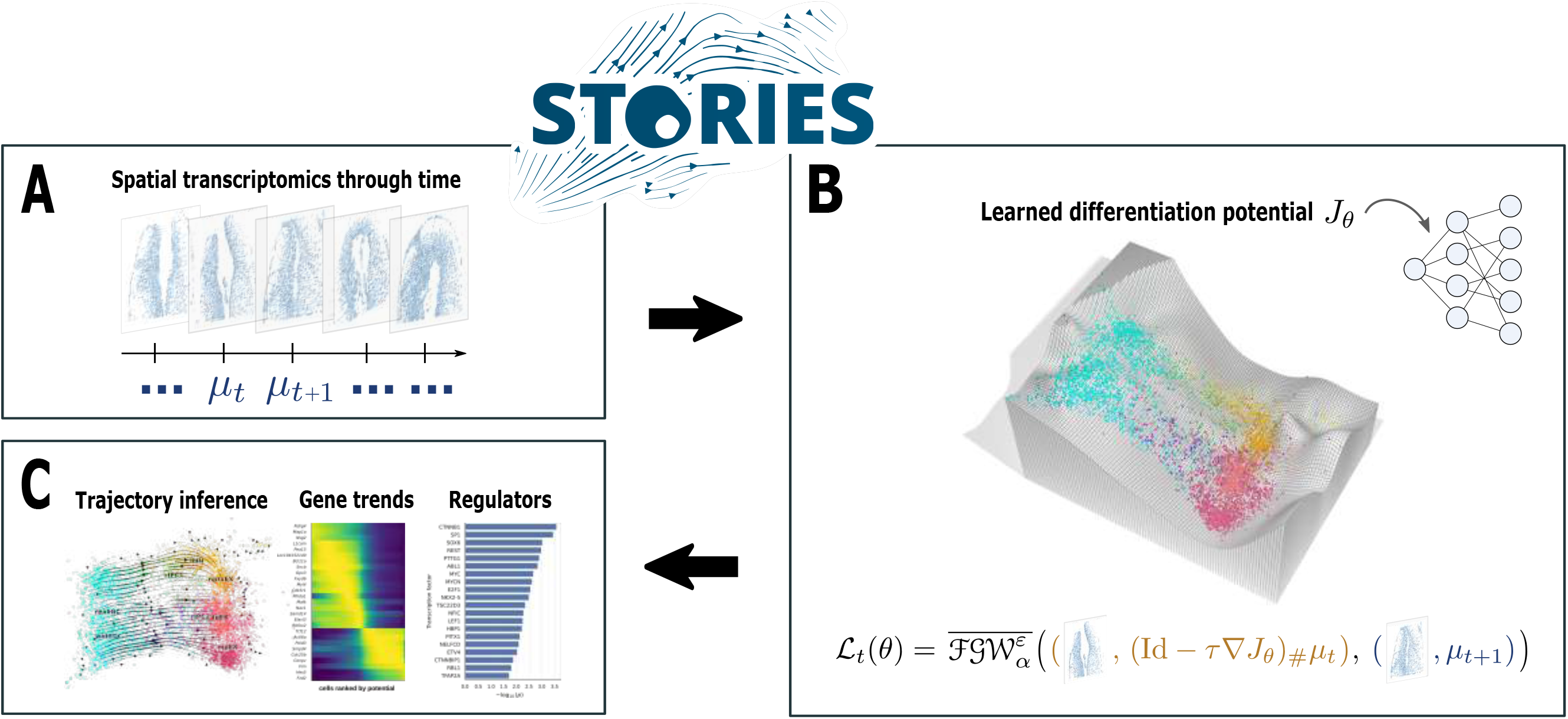
Overview of STORIES. **(A)** STORIES takes as an input spatial transcriptomics through time. **(B)** STORIES learns the parameters *θ* of a neural network *J*_*θ*_ representing the differentiation potential of a cell based on its transcriptomic profile. The objective function is based on Fused Gromov-Wasserstein, which leverages both the transcriptomic profile and the spatial coordinates. **(C)** The gradient of the function *J*_*θ*_ delivers a velocity that can be used to perform trajectory inference. The potential itself is a natural alternative to pseudotime and allows the study of gene trends along differentiation. Finally, STORIES can highlight possible transcription factors regulating differentiation.

STORIES is based on the Optimal Transport (OT), a mathematical framework that enables the geometrically meaningful comparison of distributions, using various flavors of the Wasserstein distance^29^. OT also provides a valuable model for population dynamics: the so-called Wasserstein gradient flows were popularized by Jordan, Kinderlehrer, and Otto for their connection with the Fokker-Planck equation and were recently used for trajectory inference in single-cell transcriptomics^15–17,30^. However, existing methods for trajectory inference based on Wasserstein gradient flows are not equipped to deal with spatially resolved omics data. STORIES introduces key methodological innovations that allow one to address the specific challenges of including spatial information.

As an input, STORIES takes slices of spatial transcriptomics profiled at several time points. For instance, Figure 1A displays sections of axolotl brains profiled at different stages during regeneration. STORIES then learns the parameters *θ* of a neural network *J*_*θ*_, which assigns a differentiation potential to each cell according to its gene expression profile *x* (see Figure 1B). Of note, this potential is not a function of space but only of gene expression. The function *J*_*θ*_ formalizes the Waddington epigenetic landscape, where undifferentiated cells have a high potential and, as they differentiate, move towards low-potential transcriptomic states, which correspond to mature cell types^18^. The transition to these low-potential attractor states defines a causal model of cellular dynamics capable of predicting future gene expression patterns and suggesting potential driver genes and transcriptional regulators (see Figure 1C).

STORIES’s potential-based approach provides two interpretable and biologically meaningful outputs: (i) the potential *J*_*θ*_ (***x***), naturally orders cells ***x*** along a differentiation process (ii) the vector - ∇_*x*_ *J*_*θ*_ (***x***) gives the direction of the evolution of gene expression. On the contrary, pseudotime-based methods^9,31^ focus on the first aspect, and velocity-based^11,32^ methods focus on the second. Crucially, STORIES also innovates compared to state-of-the-art potential-based methods^15–17^ by enabling the use of spatial coordinates.

Briefly, STORIES trains the neural network *J*_*θ*_ by predicting a distribution 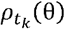 of gene expression profiles for each time point *t*_*k*_ where *k* ∈ [1,…, *K*]. These distributions are then compared to the ground-truth distributions 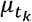, and the parameters *θ* are updated to improve the predictions. Unlike existing potential-based methods, STORIES allows one to take into account the spatial coordinates of cells when comparing the distributions of gene expression.

Formally, let 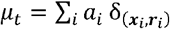, denote the empirical distribution of cells at time *t*, characterized by their gene expression profile ***x***_*i*_ ∈ ℝ^*d*^, spatial coordinates ***r***_*i*_ ∈ ℝ^*2*^ and weight *a*_*i*_ ∈ ℝ_+_ where ∑_*i*_*a*_*i*_= 1. Similarly, let us denote 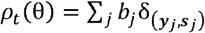, ? the predictions of STORIES at time *t*. Unlike the gene expression profiles ***x***_*i*_, ***Y***_*j*_, the spatial coordinates ***r***_*i*_, ***s***_*j*_ are not directly comparable because the slices are not necessarily aligned between time points. In other words, the spatial coordinates ***r***_*i*_, ***s***_*j*_ are defined up to an isometry (e.g., a rotation or translation).

Existing potential-based methods train the neural network using a linear OT objective, which is sensitive to isometries. Our approach instead uses a recently developed quadratic extension of OT called Fused Gromov-Wasserstein (FGW)^23^, which renders the model invariant to spatial isometries. The FGW distance, defined below and explained more thoroughly in the Methods section, allows one to compare the distributions *μ*_*t*_ and *ρ*_*t*_ directly on gene expression profiles, and up to an isometry on spatial coordinates.

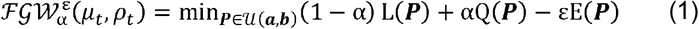

FGW seeks a matrix ***P*** mapping cells from *μ*_*t*_ to *ρ*_*t*_ such that ***P*** minimizes the sum of three terms: (i) the linear term L compares the gene expression coordinates ***x***_*i*_, ***Y***_*j*_ (ii) the quadratic term *Q* compares pairwise distances 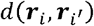 and 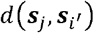, which are not affected by translating or rotating the tissue (iii) an entropic regularization term E. The parameter α ∈ [0,1] denotes the relative weight of spatial information.

Our proposed objective function evaluates the predictions across all time points using a debiased version of the FGW distance denoted 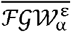 (see Methods):

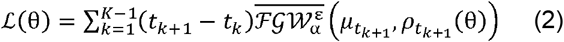

For α = 0, Equation 2 corresponds to a model relying purely on linear OT and which does not leverage spatial information, as proposed in the state-of-the-art^15–17^. In the following, we refer to this as the linear method. Existing methods^15–17^ propose different strategies to make the predictions 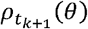. They vary in terms of teacher-forcing, number of steps between *t*_*k*_ and *t*_*k* + 1_, and whether steps are implicit or explicit (see “Discretization” in Methods). As explained in the following section, we thus compared those implementation choices in order to achieve the best performances for both STORIES and the linear method, see Extended Data Figure 1.

STORIES is implemented as an open-source Python package seamlessly integrated into the classical Python single-cell analysis pipeline (github.com/cantinilab/stories). Users can thus take advantage of scverse tools like Scanpy, Squidpy, and CellRank for preprocessing and downstream analysis^25,26,33^. In addition, STORIES provides a user-friendly visualization of driver genes and enriched transcription factors, thus helping biological interpretability. Finally, STORIES leverages gpu acceleration to train models efficiently (less than 20 minutes on a dataset of 396k cells and seven time points with an A40 Nvidia GPU).

In the following sections, we extensively benchmark STORIES against the available state-of-the-art using large-scale spatiotemporal atlases. While stVCR^34^ is the only existing method using space to perform trajectory inference, no code implementation is provided to compare it with STORIES. Regarding state-of-the-art OT-based methods which do not use spatial information, most of them don’t have a code implementation allowing their application on new data^14,17^. PRESCIENT^16^ is thus the only state-of-the-art method that could be compared to STORIES on our benchmark datasets.

### Benchmarking STORIES against the state of the art

We assessed the effectiveness of STORIES in predicting cell states over time across three Stereo-seq spatiotemporal atlases: a mouse development atlas, a zebrafish development atlas, and an axolotl brain regeneration atlas^4,6,7^. Details on data processing are provided in the Methods section.

From each atlas, we created three sets: a training set, an early test set, and a late test set (see Figure 2A). The test sets are composed of two time points, and the goal is to use the first time point to predict the second time point’s gene expression. The late test set is particularly challenging because its second slice comes from an entirely new time point, which may contain cell states not seen during training. For example, in the zebrafish atlas, fast muscle cells only appear at 24 hours post-fertilization (i.e. hpf), whereas the training set includes slices only up to 18 hpf.

**Figure 2.**
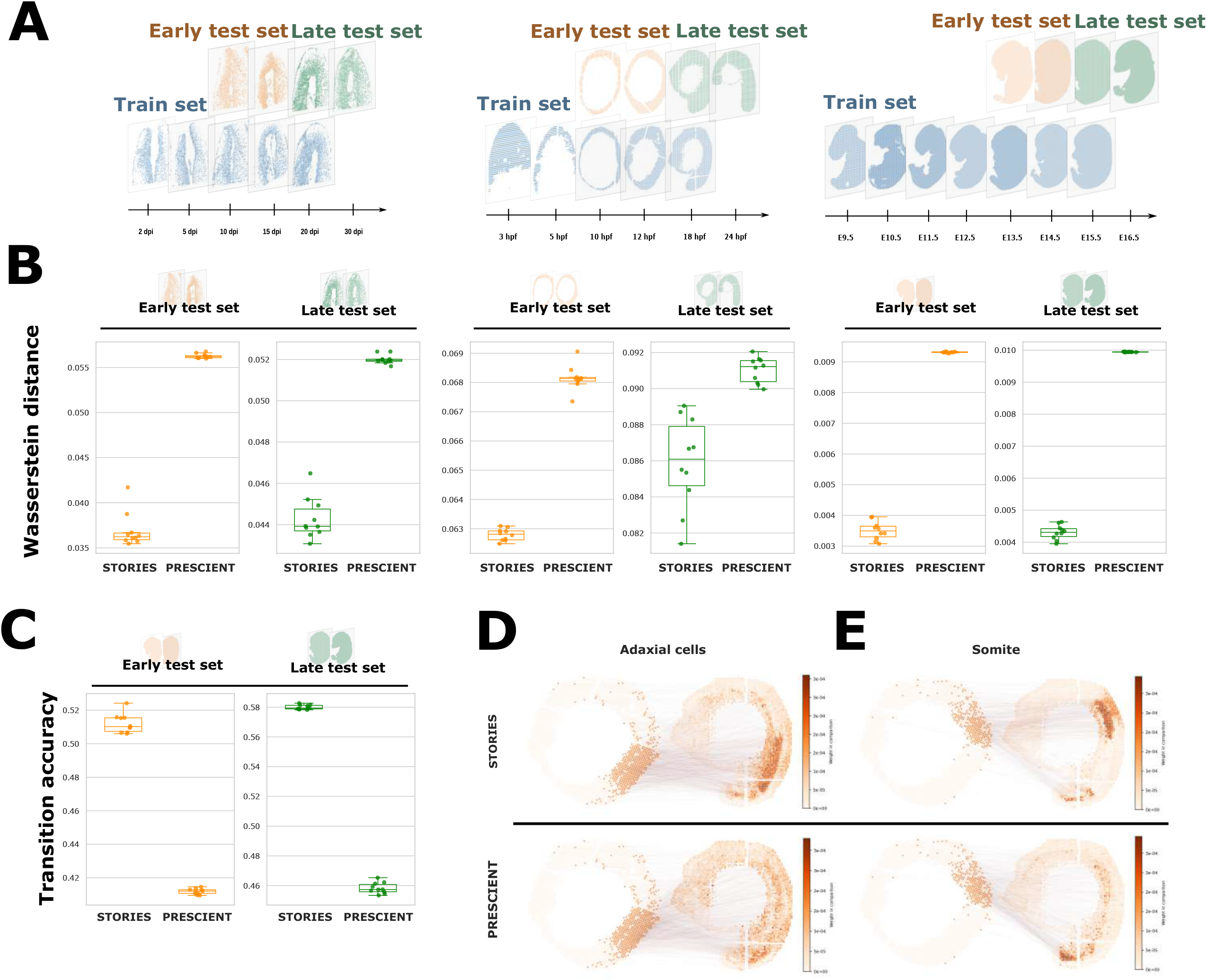
Benchmark of STORIES on three large datasets. **(A)** Visual representation of the three datasets in our benchmark. From left to right, an axolotl brain regeneration dataset, zebrafish development, and mouse development. Slices in each dataset are split into a train set (blue), an early test set (orange), and a late test set (green). **(B)** Wasserstein distance between predicted and ground truth gene expression in early test sets (orange) and late test sets (green) across the three datasets. Scores are reported for n = 10 initialization seeds. In the boxplots, the center line, box limits, and whiskers denote the median, upper and lower quartiles, and 1.5× interquartile range, respectively. **(C)** Cell type transition accuracy on the mouse development dataset in the early test set and late test set. Scores are reported for n = 10 initialization seeds. In the boxplots, the center line, box limits, and whiskers denote the median, upper and lower quartiles, and 1.5× interquartile range, respectively. **(D)** Visual representation of the linear optimal transport matching between the gene expression predicted by STORIES (top) or PRESCIENT (bottom) and the target gene expression measured at the next time point for two specific cell populations of the zebrafish development dataset, Adaxial cells (left) and Somite (right). For each example, the left slice displays the positions of cells belonging to that cell population and the right slice displays the cells they are matched with at the following time point.

*α* is a key parameter in STORIES as it weighs the relative importance of the spatial information in the FGW loss. In order to set a value for *α* in the benchmark we want to obtain the best tradeoff between accurately predicting gene expression at the slice level and making a prediction that is consistent with space. To optimally evaluate this tradeoff we solve a FGW problem between STORIES’ gene expression predictions and the ground truth measurements and use the linear term of the solution as a gene expression error and the quadratic term as a measure of spatial consistency (see Methods). Based on this evaluation, (Extended Data Figure 2A) setting an *α* in the range [1x 10^−3^,…,5 × 10^−3^] achieves the best tradeoff. We therefore use α = 5 × 10^−3^ for the rest of the benchmark.

To evaluate the impact of spatial information, we compared STORIES with its linear counterpart *α* = 0). Since the latter incorporates the best-performing aspects of state-of-the-art methods^15–17^ for our experiments (see “Discretization” in Methods and Extended Data Figure 1) and shares the same backbone as STORIES, it offers an unbiased way to assess the usefulness of space in trajectory inference. For this comparison, we used the classical metric in the context of trajectory inference: the Wasserstein distance^29^. This metric is only used for evaluation since it disregards the spatial information and thus would not allow us to visualize the tradeoff between spatial consistency and gene expression prediction. As shown in Extended Data Figure 2B, adding the space improves performance according to the Wasserstein metric in all test cases. Additionally, we displayed in Extended Data Figure 2C-E qualitative examples showing why the training loss of STORIES is more biologically relevant than its linear counterpart. Since both methods involve matching predictions with a reference population of cells, we compared their matchings for specific cell types (see Methods). First, in the axolotl atlas (Extended Data Figure 2C), STORIES correctly matches predictions from immature neurons (IMN) with Nptx+ excitatory neurons in the lateral pallium (NptxEX), and predictions from regeneration intermediate progenitor cells (rIPC2) with excitatory neurons in the dorsal pallium (dpEX)^7^. The linear method, on the contrary, incorrectly matches rIPC2 predictions with microglial cells (MCG) from a different anatomical region. Second, in the zebrafish atlas (Extended Data Figure 2D), STORIES accurately matches predictions from the optic vesicle with cells located around the eye, and predictions from the polster with cells located within the head. In contrast, the linear method incorrectly matches optic vesicle predictions with a broad group of cells across different anatomical regions, and polster predictions with cells from the tail area. Third, in the mouse atlas (Extended Data Figure 2E), STORIES correctly matches predictions from liver and lung cells with their respective organs. The linear method, instead, incorrectly matches lung cell predictions with a broad group of cells across organs.

We now benchmark STORIES against PRESCIENT which is the only available state-of-the-art method for the benchmark datasets, see Methods. As shown in Figure 2B, according to the Wasserstein distance, STORIES outperforms PRESCIENT on all test cases. We then evaluated the cell type transition accuracy of both approaches on the mouse dataset, see Figure 2C. Indeed, for mouse development, biologically valid cell type transitions were provided by Klein D. et al^21^. We can thus assess to which extent the predictions of STORIES and PRESCIENT fit with this ground truth, see Methods. STORIES outperforms PRESCIENT also according to this evaluation metric.

Finally, in Figure 2D-E we display practical examples where STORIES, due to its innovative usage of the space, identifies better trajectories than PRESCIENT (see Methods). In zebrafish development when transitioning from 12HPF to 18HPF, PRESCIENT wrongly predicts Adaxial cells to be distributed all over the embryo (Figure 2D). On the contrary, STORIES correctly matches Adaxial cells with cells close to the notochord. Indeed, adaxial cells are known to reside next to the notochord^35^. In addition, Adaxial cells are known to differentiate into slow muscle cells^35^ and indeed based on the annotation of 24HPF (see Extended Data Figure 3A), slow muscle cells are localized in the same area where STORIES predicts Adaxial cells to evolve. In addition, in the same setting, PRESCIENT wrongly predicts somite cells to match only with the tail area missing to identify most of the somite cells annotated in the 18HPF zebrafish embryo (Figure 2E). On the opposite, STORIES correctly matches somite cells from 12HPF with all somite cells from 18HPF. See Extended Data Figures 3B-C for other cell types in the zebrafish dataset. Finally, also in axolotl transition from 15DPI to 20DPI, we can observe a stronger mapping for wntEGCs and RrIPC1 cells in STORIES (Extended Data Figure 4A) with respect to PRESCIENT (Extended Data Figure 4B). In the same way, in mouse transition from E14.5 to E15.5, we observe a more accurate mapping for “Choroid plexus” and “jaw and tooth” in STORIES (Extended Data Figure 5) with respect to PRESCIENT (Extended Data Figure 6).

STORIES’s superior performance in achieving biologically coherent and accurate gene expression predictions demonstrates the significant benefit of considering spatial information when learning a gradient flow model on spatial transcriptomics data.

### STORIES reveals drivers of neuron regeneration in axolotls

To further assess the potential of STORIES for trajectory inference in spatial transcriptomics through time, we first focused on axolotl brain regeneration.

We trained STORIES as described in Methods on the subset of cells described in the original publication as involved in neuron regeneration: Wnt+ and reactive ependymoglial cells (wntEGC and reaEGC), regeneration intermediate progenitor cells (rIPC1 and rIPC2), immature neurons (IMN), Nptx+ lateral pallium excitatory neurons (nptxEX), dorsal pallium excitatory neurons (dpEX), and medial pallium excitatory neurons (mpEX)^7^. As shown in Figure. 3A, STORIES learns an energy landscape consistent with the original publication. Indeed, the potential *J*_*θ*_ assigns a high potential to progenitor states (wntEGC and reaEGC), a medium potential to intermediary states (rIPC1, rIPC2, and IMN), and a low potential to mature states (nptxEX, dpEX, and mpEX).

**Figure 3.**
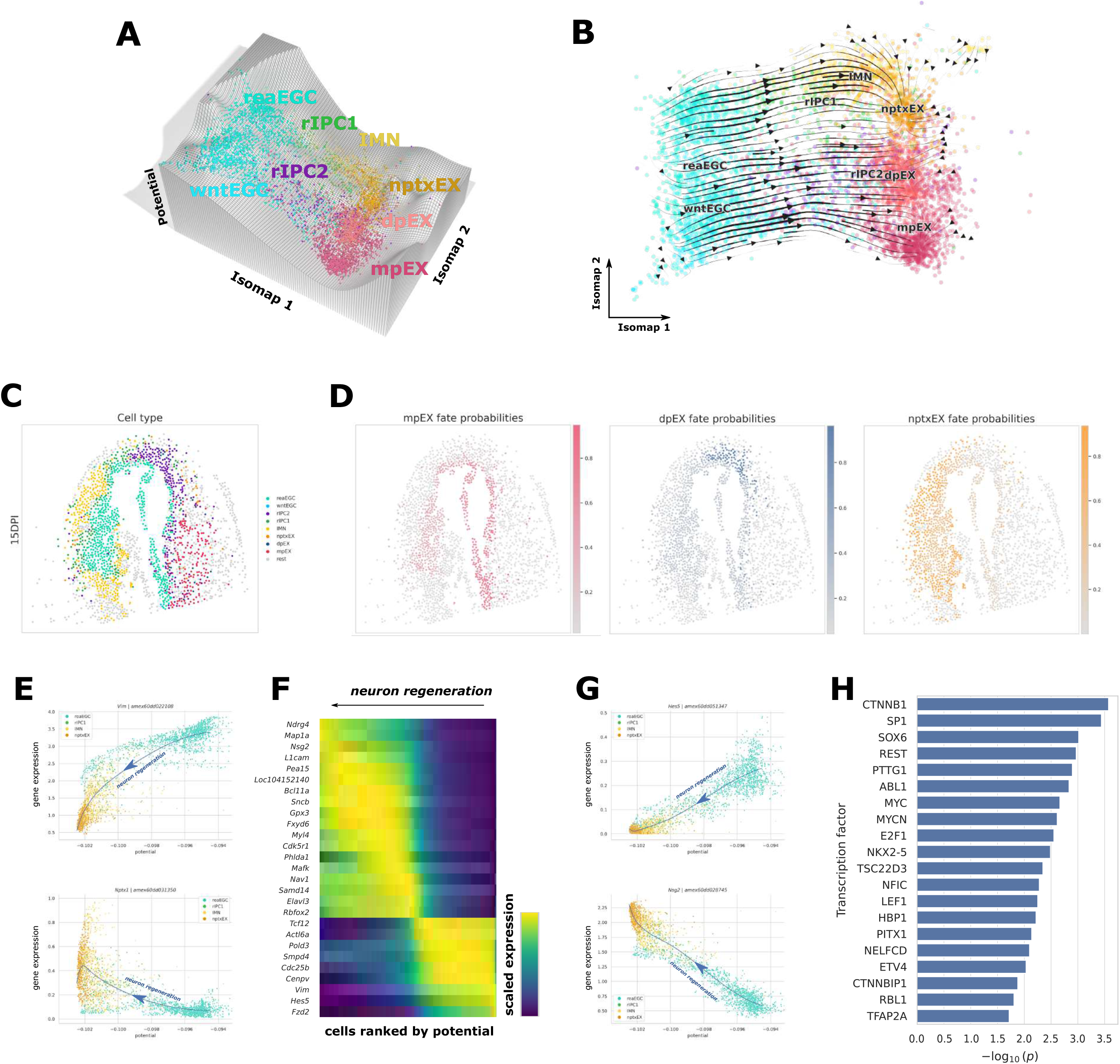
Trajectory inference with STORIES in axolotl neuron regeneration. **(A)** 3-D representation of the potential landscape learned with STORIES. The *x* and *y* axes are Isomap coordinates, and the *z*-axis is an interpolation of the potential. Colors represent cell types involved in the regeneration process. **(B)** Visual representation of cell-cell transitions computed using CellRank from STORIES’s velocity vectors. **(C)** 2-D visualization of an axolotl slice at 15DPI, with cells colored by their cell type annotation. **(D)** 2-D visualization of an axolotl slice at 15DPI, with reaEGCs colored by their predicted fate probabilities for mpEX, dpEX and npxTX fates, from left to right. **(E)** Smoothed gene expression for *Vim* and *Nptx1* along the potential computed by STORIES. The blue line is a spline regression of expression from potential. **(F)** Normalized gene expression regressed using a spline model along the potential computed by STORIES. Genes are ordered by the potential for which they achieve maximum expression **(G)** Smoothed gene expression for *Hes5* and *Nsg2* along the potential computed by STORIES. The blue line is a spline regression of expression from potential. **(H)** Enrichment scores of Transcription Factors targeting candidate driver genes. A one-sided Wilcoxon rank-sum test is used to report p-values.

We computed cell-cell transitions by applying CellRank on the gradient of the trained potential (see Methods), as visualized in Figure 3B. These transitions highlight that STORIES not only detects the correct stage of differentiation, but also recovers the three major trajectories described in the original publication: wntEGC-mpEX, reaEGC-rIPC2-dpEX, and reaEGC-rIPC1-IMN-nptxEX^7^. For an overview of cell type transitions see Extended Data Figure 7A. Importantly, the authors identified these trajectories by applying Monocle separately on three spatial regions and specifying EGCs as the starting point in each case. In contrast, STORIES achieves the same results without the need to isolate specific spatial regions and specify the starting point of the trajectory. Indeed, STORIES leverages spatial information to process all regions simultaneously and leverages temporal information to infer progenitor states from the data.

We then investigated how the spatial environment influences these cell trajectories. Focusing on 15DPI, we observed that the reaEGCs have a different cell fate depending on their spatial location in the slice (Figure 3C-D, see Extended Data Figure 7B for the other time points and Methods for how cell fate probabilities are computed). ReaEGCs on the right of the injury tend to commit more towards mpEX, while reaEGCs on the left of the injury tend to commit more towards nptxEX. This observation seems to be in accordance with the spatial location of the terminal states in the slice. In addition, this same spatial organization of the terminal states can be observed in the additional two replicates available (Extended Data Figure 7C), supporting its biological relevance. These results suggest that the spatial environment impacts cell fate decisions. As a consequence, STORIES, integrating the spatial context, is a powerful tool for cell fate inference.

We then narrowed further into the reaEGC-rIPC1-IMN-nptxEX trajectory, which the original publication studies in most detail^7^. First, we sought to confirm expected gene trends along this trajectory. The original study suggests *Vim*, which encodes a critical cytoskeletal protein, as a marker of reaEGC cells and *Nptx1*, which is involved in synaptic plasticity, as a marker of NptxEX. Accordingly, STORIES recovered a clear decreasing trend for *Vim* expression along differentiation and a clear increasing trend for *Nptx1* expression (see Figure 3E).

Next, we performed unsupervised discovery of gene trends by fitting a spline regression model along the previously mentioned trajectory (see Methods). Figure 3F reports the best candidate driver genes across differentiation stages. Interestingly, the early stages of differentiation coincide with high expression of *Hes5* (Figure 3G), which is known to maintain stemness in the context of neural differentiation^36^, and *Cdc25b*, a cell-cycle regulator key to neuron production^37,38^. Conversely, late stages of differentiation coincide with high expression of the microtubule-associated protein gene *Map1a*, crucial to neural development and regeneration^39^, and *L1cam*, shown to promote axonal regeneration^40^. STORIES also outputs additional genes that represent possible drivers of neuron regeneration and would require further biological investigation. For instance, STORIES uncovered a trend for late expression of the scarcely studied *Nsg2* (Figure 3G), which is thought to be involved in synaptic function and, like *Nptx1*, interacts with AMPA receptors^41^.

Finally, our analysis revealed possible transcriptional regulators of the differentiation process (see Figure 3H) by testing transcription factor (TF) enrichment using the curated literature-based TRRUST database (see Methods). The most significantly enriched TF, CTNNB1, encodes β-catenin, which has been described as an essential regulator in neuron regeneration in mouse models and in limb regeneration in axolotl^42–44^. Other top TFs include SP1 and MYC, described in the context of neuron regeneration and computationally retrieved in axolotl limb regeneration^45–48^. Additionally, we identify SOX6, MYCN, and REST, which are not widely studied in the context of regeneration but are known regulators in development^49–51^. Interestingly, a recent study predicted REST as a regulator of neuron regeneration and validated this role in a mouse model^52^.

Thus not only can STORIES learn a Waddington landscape that implicitly captures the impact of space on neuron regeneration in axolotls, but it also recovers its underlying regulatory landscape through the unbiased discovery of potential drivers and mechanisms, possibly relevant for further biological investigations.

### STORIES identifies drivers of gliogenesis in mouse midbrain

We then sought to highlight STORIES’s potential in trajectory inference by studying mouse dorsal midbrain development.

We trained STORIES as described in Methods on the subset of cells described in the original article as exhibiting a branching trajectory: radial glial cells (RGC) differentiating into either neuroblasts (NeuB) or glioblasts (GlioB)^4^. As visualized in Figure 4A, STORIES learns an energy landscape consistent with the original publication. Indeed, the potential *J*_*θ*_ assigns a high potential to RGC and a low potential to the more differentiated NeuB and GlioB.

**Figure 4.**
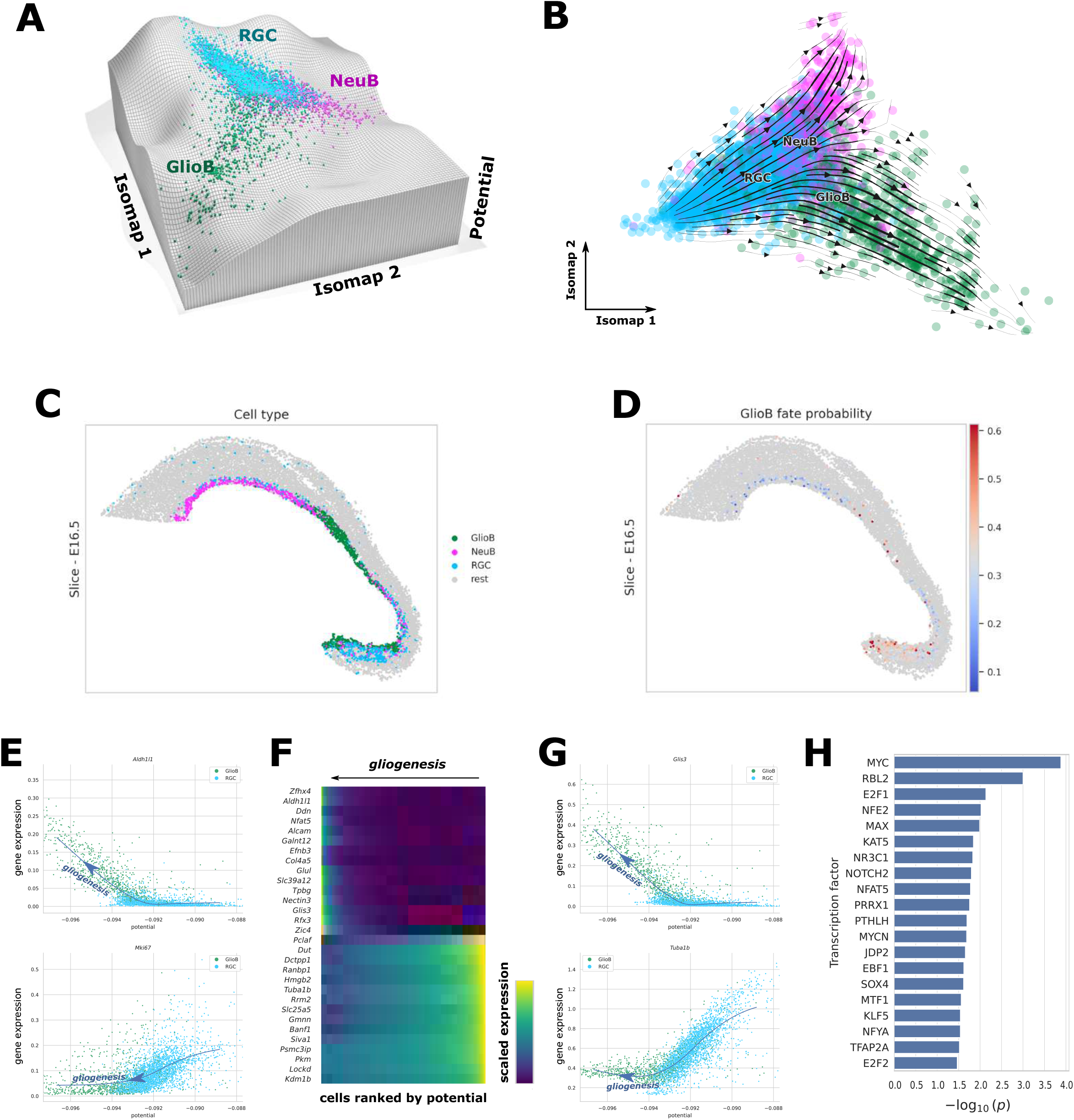
Trajectory inference with STORIES in mouse gliogenesis. **(A)** 3-D representation of the potential landscape learned with STORIES. The *x* and *y* axes are Isomap coordinates, and the *z-axis* is an interpolation of the potential. Colors represent radial glial cells which differentiate into either neuroblasts or glioblasts. **(B)** Visual representation of cell-cell transitions computed using CellRank from STORIES’s velocity vectors. **(C)** 2-D visualization of a mouse slice at E16.5, with cells colored by their cell type annotation. **(D)** 2-D visualization of a mouse midbrain slice at E16.5, with RGCs colored by their predicted probability of becoming GlioBs. **(E)** Smoothed gene expression for *Mki67* and *Aldh1l1* along the potential computed by STORIES. The blue line is a spline regression of expression from potential. **(F)** Normalized gene expression regressed using a spline model along the potential computed by STORIES. Genes are ordered by the potential for which they achieve maximum expression **(G)** Smoothed gene expression for *Tuba1b* and *Glis3* along the potential computed by STORIES. The blue line is a spline regression of expression from potential. **(H)** Enrichment scores of transcription factors targeting candidate driver genes. A one-sided Wilcoxon rank-sum test is used to report p-values.

We computed cell-cell transitions by applying CellRank on the gradient of the trained potential (see Methods), as visualized in Figure 4B. These transitions highlight that STORIES not only detects the correct stage of differentiation but also recovers the expected branching from RGC to glial and neural cell fates^4^. Importantly, the original publication identified this branching using Monocle 3, which required manually setting RGC as the trajectories’ starting point. On the contrary, STORIES achieves the same results without manual input by leveraging temporal information to infer the starting point from the data.

We then investigated how the spatial environment influences these cell trajectories. Focusing on E16.5, the differentiation of RGCs into either NeuB or GlioB seems to be influenced by their spatial location (Figure 4C-D, see Extended Data Figure 8A for the other time points and Methods for how cell fate probabilities are computed). RGCs on the rostral part tend to commit to NeuB, RGCs on the extreme side of the caudal part tend to commit to GlioB. Finally, RGCs on the central part tend to be organized into clusters of cells that either commit to NeuB or to GlioB. Importantly, these conclusions are supported by the agreement between the spatial position of the already differentiated cells and our predicted fate probabilities for the RGCs. These same observations apply for time point E14.5 (see Extended Data Figure 8A). In addition, this same spatial organization of the terminal states can be observed in the additional two replicates available (Extended Data Figure 8B), supporting its biological relevance. These results suggest that the spatial environment impacts cell fate decisions. As a consequence, STORIES, integrating the spatial context, is a powerful tool for cell fate inference.

Glial cells outnumber neurons in the brain, but their development has been studied less extensively^53^. Moreover, understanding gliogenesis is of critical therapeutic importance because of its parallels with glioma, the most common and deadliest form of brain cancer^54^. Thus, we focused further on the RGC-GlioB trajectory. We first sought to confirm expected gene trends along this trajectory. The original study identifies *Mki67*, a proliferation marker, as highly expressed in RGC, and *Aldh1l1*, an astrocyte marker, as highly expressed in GlioB^4^. Accordingly, STORIES recovered a decreasing trend for *Mki67* expression along differentiation and an increasing trend for *Aldh1l1* expression (see Figure 4C).

Next, we performed unsupervised discovery of gene trends by fitting a spline regression model along the previously mentioned trajectory (see Methods). Figure 4D reports the best candidate driver genes across differentiation stages. The early stages of differentiation coincide with a high expression of cell cycle genes *Gmnn, Rrm2*, and *Hmgb2*^55^. Additionally, we observed a high expression of the alpha-tubulin gene *Tuba1b* in the early stages of differentiation, as previously described in the developing brain^56^. Conversely, the late stages of differentiation coincide with the high expression of the glutamine synthetase gene *Glul*, a key astrocyte marker^57,58^. STORIES also outputs additional genes that represent possible drivers of gliogenesis and would require further biological investigation. For instance, *Glis3* displays an increasing trend along gliogenesis (see Figure 4E) but is little studied in this context. However, *Glis3* was recently suggested as a therapeutic target to suppress proliferation in glioma^59^.

Finally, our analysis revealed candidate transcriptional regulators of the differentiation process (see Figure 4F) by testing transcription factor (TF) enrichment using the curated literature-based TRRUST database (see Methods). Among the most enriched TFs, SOX4 and NOTCH2 have been studied in gliogenesis^60,61^. Additionally, STORIES recovers MYC, MYCN, and MAX, which have been studied in the context of glioma^62^.

This second experiment confirms that not only STORIES learns a Waddington landscape that implicitly captures the impact of space on gliogenesis in mice, but it also recovers its underlying regulatory landscape through the unbiased discovery of potential drivers and mechanisms that motivate further biological investigations.

## Discussion

Recent technological advances in spatial transcriptomics have enabled the tracking of gene expression at single-cell resolution in the spatial context of the tissue. Large datasets of spatial transcriptomics profiled through time give a unique opportunity to understand dynamic biological processes such as development and disease onset. However, their analysis requires trajectory inference tools tailored to the specific challenges of spatial data.

In this article, we proposed SpatioTemporal Omics eneRgIES (STORIES), a computational framework for trajectory inference from spatial transcriptomics profiled at several time points. STORIES enables a rich and spatially informed analysis of differentiation trajectories. To evaluate STORIES’s performance, we benchmarked it against the state-of-the-art in three large Stereo-seq datasets and highlighted the advantage of considering spatial information in trajectory inference from time-course single-cell data. We further showcased STORIES’s abilities in two concrete settings: axolotl neuron regeneration and mouse gliogenesis.

STORIES offers a model of population dynamics tailored for single-cell resolution spatial transcriptomics technologies like Stereo-seq or Visium HD. Given the fast-paced developments in spatial transcriptomics^1^, the number of spatiotemporal atlases at single-cell resolution can be expected to increase steadily. At the same time, STORIES could be applied to low-resolution data (e.g. 10x Visium), which have a spot size larger than the typical cell, using deconvolution techniques^63^. In addition, STORIES could be adapted to imaging-based technologies like MERFISH which offer high resolution but can only detect a limited panel of genes^64^.

STORIES provides an interpretable model of differentiation relying on a potential energy. Previous work shows that such potential landscapes arise naturally from simple gene regulatory networks (GRNs)^19^. However, potential energies cannot model complex GRNs, cell-cell communication, or oscillations within a cell state^65^. Extensions to more complex energy functionals could thus lead to further insights into biological processes such as development or immune response. For instance, although numerically challenging because of their quadratic evaluation cost, interaction energies represent a very exciting opportunity to integrate cell-cell communication in trajectory inference models^66^ in order to study the onset of complex diseases.

The major novelty of STORIES is its ability to learn a spatially-informed potential. This methodological development is critical because dynamic processes such as development involve coordinated expression changes and tissue reorganization^67^. However, the learned potential operates only on gene expression, so it does not allow the prediction of future positions of cells. Including a spatial component in the energy function may provide a more comprehensive view of biological processes by predicting cell migration. Combining such an energy with the Gromov Wasserstein geometry could help define more complex flows. Despite recent works^68^, the extension of Wasserstein flows to Gromov-Wasserstein flows remains poorly understood theoretically since its introduction by Sturm^69^ and is currently infeasible for practical implementation. Overcoming these challenges would pave the way for a more comprehensive modeling of biological processes. Relating this to existing models for morphogenesis, such as Alan Turing’s reaction-diffusion model^70^, is an exciting avenue for further research.

## Supporting information

Supelementary text

## Acknowledgements

The project leading to this manuscript has received funding from the European Union, ERC StG, MULTIview-CELL, 101115618 (L.C.). In addition, this work has been funded by the French government under management of Agence Nationale de la Recherche as part of the “Investissements d’avenir” program, reference ANR-19-P3IA-0001; PRAIRIE 3IA Institute (L.C. and G.P.) and by the Inception program “Investissement d’Avenir grant ANR-16-CONV-0005” (L.C.). The work of G. P. was supported by the European Research Council (ERC project NORIA). We acknowledge the help of the HPC Core Facility of the Institut Pasteur and Déborah Philipps for the administrative support.

## Author Contributions Statement

G-J.H., G.P. and L.C. designed and planned the study. G-J.H. developed the tool. G-J.H. and J.S. performed most analyses. A.A. contributed to the growth rate analysis. D.C. helped with discussing the impact of space on cell fate trajectories. G-J.H., J.S. and L.C. wrote the paper. G.P. revised the manuscript. All authors read and approved the final manuscript.

## Methods

### Data collection

#### Zebrafish

For the training and validation sets, we used the five first time points showcased in Figure 2A of the original publication^6^: 3.3hpf slice 1, 5.25hpf slice 10, 10hpf slice 11, 12hpf slice 8, 18hpf slice 8.

For the test set, we used:

- 10hpf slice 17 and 12hpf slice 5 to evaluate prediction within the time range seen during training. These two slices were studied together in Figure S4A of the original publication^6^.
- 18hpf slice 11 and 24hpf slice 4 to evaluate prediction outside the time range seen during training. These two slices were studied together in Figure S4E of the original publication^6^.

Altogether, this represented 17,920 cells after preprocessing.

#### Mouse

For the training and validation sets, we used the 7 first time points showcased in Figure 3A of the original publication^4^: E9.5 E1S1, E10.5 E1S1, E11.5 E1S1, E12.5 E1S1, E13.5 E1S1, E14.5 E1S1, E15.5 E1S1.

For the test set, we used:

- E13.5 E1S2 and E14.5 E1S2 to evaluate prediction within the time range seen during training. These two slices were studied in Figure S2C of the original publication^4^.
- E15.5 E1S2 and E16.5 E1S1 to evaluate prediction outside of the time range seen during training. The first slice was studied in Figure S2C and the second in Figure 3A of the original publication^4^.

Altogether, this represented 794,063 cells after preprocessing.

#### Dorsal midbrain

We retained one slice per time point. We used the 3 slices showcased in Figure S7A of the original publication^4^: E12.5 (E1S3), E14.5 (E1S3), E16.5 (E1S3). As in the original publication, we subset the analysis to the RGC, NeuB, and GlioB cell types.

Altogether, this represented 4,581 cells after preprocessing. Note that the “Mouse” dataset and the “Dorsal midbrain” dataset originate from the same experiments but have different resolutions (bin 50 vs image-based segmentation), so the same neural network weights cannot be used in both cases.

#### Axolotl

*Benchmark*. For the training and validation sets, we used the five first time points showcased in Figure 3B of the original publication^7^: 2DPI (rep1), 5DPI (rep1), 10DPI (rep1), 15DPI (rep3), 20DPI (rep2). We manually removed spatial outliers in 10DPI (rep1), 15DPI (rep4), and 20DPI (rep2). For the test set, we used:

- 10DPI (rep2) and 15DPI (rep1) to evaluate prediction within the time range seen during training. These two slices were showcased in Figure S9 of the original publication^7^.
- 20DPI (rep3) and 30DPI (rep2) to evaluate prediction outside of the time range seen during training. These two slices were showcased in Figure S9 of the original publication^7^.

As in the original publication, we restricted the analysis to the dorsal part of the injured hemisphere. Altogether, this represented 22,083 cells.

#### In-depth analysis

For the analysis in Section 3 of Results, we used slices 2DPI (rep1), 5DPI (rep1), 10DPI (rep1), 15DPI (rep3), 20DPI (rep2), and 30DPI (rep2). As before, we manually removed spatial outliers and restricted the analysis to the dorsal part of the injured hemisphere. As in the original publication, we subset the data to the following cell types: nptxEX, reaEGC, wntEGC, dpEX, mpEX, IMN, rIPC1, rIPC2. This represented 5,904 cells.

### Preprocessing

All datasets were stored and handled as Numpy arrays^71^ wrapped in Anndata objects^72^. For all datasets, we performed the following preprocessing steps.

#### Cell and gene quality control

Using Scanpy’s sc.pp.filter_cells, we removed cells with less than 200 expressed genes. Then, we removed the top 0.1% of cells with the most expressed genes. Finally, we removed genes expressed in less than 3 cells using Scanpy’s sc.pp.filter_genes.

#### Normalization and highly variable gene selection

We applied Scanpy’s “Pearson residuals normalization” and selected 10,000 highly variable genes. Scanpy computed highly variable genes for each batch and merged them to avoid selecting batch-specific genes.

#### Dimensionality reduction

Using Scanpy’s sc.tl.pca, we applied Principal Component Analysis (PCA) to reduce the data to 50 dimensions. Section 3 and Section 4 of Results used a subset of relevant cell types. This was done after PCA but before batch correction.

#### Batch correction

We applied Harmony on the PCA components to correct the batch effects, using Scanpy’s sc.external.pp.harmony_integrate, a wrapper around harmonypy^73^.

#### Visualization

Using Scanpy’s sc.tl.umap, we applied UMAP to project the batch-corrected data into two dimensions. In Section 3 and Section 4 of Results we applied Isomap instead of UMAP. Indeed, we found that visually, Isomap respected cell type transitions better than UMAP. We used scikit-learn’s sklearn.manifold.Isomap.

### Wasserstein gradient flow learning with a quadratic objective Notations

Let us consider a time point *t* ∈ ℝ and *μ*_*t*_ ∈ 𝒫(ℝ)^*d*^ a discrete distribution of *n* cells.

We denote 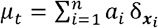 where the vector *x*_*i*_ ∈ ℝ^*d*^ represents the gene expression of the *i*-th cell, and the weights *a*_*i*_ ∈ ℝ _+_ are such that ∑ *a*_*i*_ = 1. In the following, for a function ?f: ℝ ^*d*^ → ℝ ^*d*^, we define the pushforward measure as 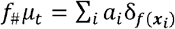.

### Wasserstein gradient flow

Similarly to previous works^15–17^, we model the evolution of *μ*_*t*_ as a Wasserstein gradient flow for a potential energy. For one single cell ***x***, the Euclidean gradient flow is a ***x***_t_ that verifies 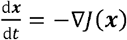, the continuous counterpart of gradient descent. Wasserstein gradient flows extend this to the space of measures^74^.

We say that *μ*_*t*_ is a Wasserstein gradient flow for the energy ℱ: µ ↦, ∑_*i*_ *J* (***x***_*i*_) d µ(***x***_*i*_) ∈ ℝif its density verifies the continuity equation

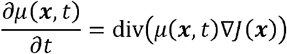

As in previous works^15–17^, we do not know the potential *J*: ℝ_*d*_ → ℝ a priori and aim instead to learn a neural network *μ*_*t*_ from snapshots 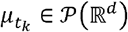 for *t*_1_ < … < *t*_*k*_ < … < *t*_*K*_. See “neural network architecture” for details about *μ*_*t*_.

### Discretization

Our approach boils down to learning *J*_*θ*_ such that for given parameters *θ* and an initial population 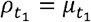, the predicted populations 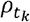 are close to the observed snapshots 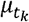. To make these predictions, existing potential-based methods^15–17^ differ in three main aspects: number of steps, teacher-forcing, and discretization scheme. As detailed below, we select the best-performing choices for these three aspects, as measured by validation loss in the Zebrafish atlas. The linear method (*α*= 0) thus differs from existing works by combining their best-performing aspects.

*Number of steps*. Hashimoto et al.^16^ and Yeo et al.^16^ make intermediary predictions between 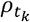 and 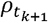, i.e. τ« (*t*_*k*+1_ - *t*_*k*)_. Bunne et al. ^18^ perform a single step instead, i.e. *τ*= (*t*_k+1_ - *t*_*k*)_^17^. In our experiments, multiple steps did not improve results (see Extended Data Figure 1A) so we chose the computationally less expensive single-step method.

#### Teacher-forcing

Hashimoto et al.^16^ and Yeo et al.^17^ predict 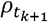 from 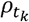. Bunne et al.^18^ introduce teacher-forcing, i.e. predicting 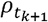 from 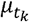 ^17^. In our experiments, teacher-forcing improved results (see Extended Data Figure 1B), so we used it throughout this work.

#### Discretization scheme

To predict 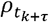 from an earlier population *ρ*_*t*_, we used the forward Euler discretization scheme ρ_*t*+*τ*_ = (Id − *τ ∇ J*_*θ*_,)_#_*ρ*_*t*_ as Hashimoto et al.^16^ and Yeo et al.^17^. In our discrete setting, this corresponds to ***x***_*t*+ τ_= ***x***_*t*_ − *τ ∇ J*_*θ (*_*x*_*t*_) for each cell ***x***. Bunne et al.^18^ propose using a backward Euler scheme to improve stability for large *τ* ^17^. We implemented this approach using the JAXopt package^75^. However we did not find this to improve results in our experiments (see Extended Data Figure 1C), so we used the computationally less expensive forward method.

### Pairwise information

Our setting differs from previous works^15–17^ in that for each cell ***x*** ∈ ℝ^*d*^ we have access to spatial coordinates ***x*** ∈ ℝ^*2*^. Spatial coordinates are defined up to an isometry. For instance, a slice of an embryo may be rotated without changing the problem. Consequently, one cannot simply concatenate ***x*** and ***r*** to leverage the spatial coordinates. The next paragraph details how we used the coordinates ***r*** to inform the problem, while still defining *J*_*θ*_:***x*** ∈ ℝ^*d*^ ↦ *J*_*θ* (_***x***) ∈ ℝ only on gene expression.

### Learning the potential

For each *k* ∈ [1,…,*K*], we compared the prediction 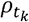 to the reference snapshot 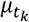.

#### Linear mode

Let us consider two discrete probability distributions 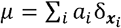 and 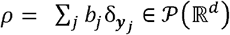. To compare *µ* and *ρ*, previous works^15–17^ use the Sinkhorn divergence^76^, defined as

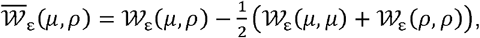

where 𝒲_ℰ_ is the entropy-regularized Optimal Transport^77^ (entropy-regularized OT), defined as

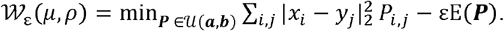

Here, 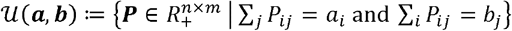 and ∑_*i*_*p*_*ij*_ = *b*_*j*_} and −∑_*i,j*_*p*_*ij* (log_ *p*_*ij*_ − _1)_ the Shannon entropy. 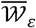 is a debiased version of 𝒲_ℰ_, such that 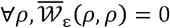.

#### Quadratic model

The Sinkhorn divergence between *ρ* and *µ* only compares distributions of gene expression. Instead, we propose a debiased Fused Gromov-Wasserstein (debiased FGW) loss to enforce the spatial coherence of the predictions. Let us consider 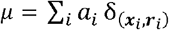, and 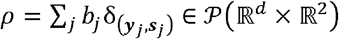. Gromov-Wasserstein (GW) is a quadratic extension of OT well suited to compare measures defined up to an isometry^78^. A debiased version of GW has been used to learn a Generative Adversarial Network (GAN)^79^. Fused Gromov-Wassertein (FGW)^23^ combines a linear and a quadratic OT term. In our setting, it is natural to use the linear term for gene expression and the quadratic term for spatial coordinates:

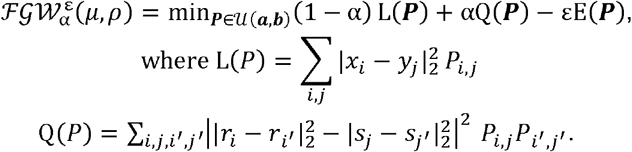

Analogously to Bunne et al.^18^, we introduce a debiased FGW to ensure the loss vanishes for an exact match^79^.

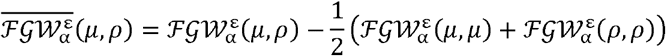

A weight α = 1 corresponds to the debiased entropy-regularized GW^80^. As α = 0, we recover the Sinkhorn divergence 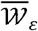.

#### Final loss function

For α ∈ [0, 1], the full objective reads

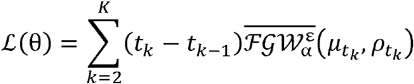

where 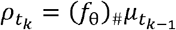 with (*f* _θ_):(***x***,***r***) ↦, (x − τ ∇ *J*_*θ*_ (***x***), ***r***). In other words, gene expression is predicted with teacher-forcing and a single forward step (see “Discretization”). Our model doesn’t predict spatial coordinates therefore ***s*** and ***r*** are identical in our FGW loss and correspond to the spatial positions of cells at time *t*_k-1_. This loss is optimized using minibatches, as done by Yeo et al.^17^ and Bunne et al.^18^ A theoretical study of minibatch Optimal Transport for debiased linear and quadratic OT is provided by Fatras et al.^81^

### Choice of the quadratic weight

To investigate the effect of the relative weight of the linear term in FGW, we reported results for α ∈ {1 × 10 ^−5^,1 × 10^− 4^,1 × 10^−3^,5 × 10^−3^,1 × 10^−2^,1 × 10^−1^}. Choosing *α*= 1 would not allow learning the potential *J*_*θ*_ because it would ignore gene expression. As expected, with a low weight (α = 10^−5^), STORIES behaves as the linear method (see Section 2 of Results).

To compare our approach with the linear model proposed in previous works^15–17^, we also trained the model with a Sinkhorn divergence, i.e. α = 0.

The benchmark in Section 2 of Results suggests good performances for values of a of the order of 10^−3^, with α = 5 × 10^−3^ performing best. The value of a can be adjusted by the user depending on the dataset. In Section 3 and Section 4 of Results we set a value of *α* =10^−3^.

### Computational Optimal Transport

#### OTT solvers

We use the OTT package to solve OT problems in a fast, GPU-enabled, and differentiable manner^28^. In particular, we rely on the Sinkhorn and GromovWasserstein solvers. We set the entropic regularization *ε* = 0.01.

#### Linear term: gene expression

The linear OT terms are defined on gene expression space, for which we chose the *d* first components of the Harmony-aligned Principal Component Analysis. We chose *d* = 50 as it is the one that maximizes cell type transition accuracy (see Evaluation section in the methods for details on how this is computed) without increasing the computational runtime of the method, see Extended Data Figure 9A-B. Notably, in both sections 3 and 4 of the results, we only used *d* = 20 as the datasets of the case studies were smaller than in the benchmark and the analysis was conducted on a subset of the cell types. Before training the neural network, we normalized the points as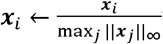 − to make the linear and quadratic terms comparable.

#### Quadratic term: Spatial coordinates

Before training the neural network and for each slice separately, we centered the spatial coordinates and scaled them to unit variance to make linear and quadratic terms comparable.

#### Neural network architecture

We implemented a Multi-Layer Perceptron (MLP) with two hidden layers of dimension 128 and GeLU activations using Flax^82^. This is a similar architecture to previous works^15–17^. The linear output layer has no bias since it would not influence the values of ∇*J*_*θ*_. Likewise, a soft activation like GeLU is preferable to the classical ReLU because we manipulate ∇*J*_*θ*_. Indeed, the derivative of ReLU is simply a unit step function, which is discontinuous and not very expressive.

### Neural network training

#### Data loading

For each time point described in “Data Collection”, 75% of the cells were used as training samples and 25% as validation. At each training or validation iteration, a batch containing 1,000 cells per time point was sampled uniformly without replacement. In development, the early time points contain fewer cells than the later time points. If less than 1,000 cells were available for a time point, we used all available cells. To reflect the train/validation split, one in four iterations performs a validation step.

### Optimizer

We used Optax’s implementation of the AdamW optimizer, with parameters b1=0.9, b2=0.999, eps=1e-8, and weight_decay=1e-4^27^. We set the learning rate using Optax’s cosine scheduler, with an initial value of 1e-2 and 10000 decay steps. To ensure convergence, when performing 10 steps, we set the learning rate to 1e-3. Similarly, when performing an implicit step, we set the learning rate to 1e-4.

#### Early stopping

We set the maximum number of iterations to 15,000 but stopped the training when the validation loss had not improved in 150 iterations. We kept the weights associated with the lowest validation loss and used the orbax package to save and load checkpoints of the model.

#### Seeds

We ran every experiment with 10 random seeds: 17158, 20181, 12409, 5360, 21712,

21781, 24802, 13630, 9668, and 651. The random seed reproducibly determines the train/validation split and weight initialization. For the plots in Figure 2C, and the analysis in Section 3 and Section 4 of Results, the experiments correspond to the randomly chosen seed 20181.

#### Computational runtime

Due to mini-batch sampling of the data samples, STORIES can scale to very large datasets. It also leverages GPU acceleration to speed up the training of the model. See Extended Data Figure 9C for a comparison of the training time of STORIES for different values of the *α* parameter on the three benchmark datasets. Overall, we observe that while using a greater *α* (like 0.1) makes the training longer due to Gromov Wasserstein being a much more computationally heavy problem than its linear counterpart, using a small alpha leads to a more than reasonable training time. Surprisingly, in some datasets, using a low alpha leads to an even faster training than using *α* = 0. While this result was not expected, we could assume that for small alphas the regularization induced by the quadratic term in FGW makes the problem converge faster as it introduces more constraint on the solution. Training progress can be monitored using the tqdm library, which provides real-time progress bars for iterative processes.

### Running PRESCIENT

We compare STORIES to PRESCIENT which also uses Optimal transport to learn a gene expression potential governing a causal model of differentiation. We used their open-source implementation (https://github.com/gifford-lab/prescient) and applied PRESCIENT with all default parameters except for *train_sd*. Indeed the default value of 0.5 for this parameter which controls the strength of the random noise in their differentiation model was too high compared to the scale of the gene expression data in the benchmark datasets. We therefore tuned this parameter for a fair evaluation and used 0.01 for *train_sd*.

### Evaluation

For the problem of choosing the optimal *α* parameter in Extended Data Figure 2A-B, we solve the FGW problem with α = 10^−3^ and ε = 10^−3^ and report the terms *L*(***P***) and *Q*(***P***) separately. Since *L*(***P***) quantifies the error in terms of gene expression, we call this quantity the *gene expression prediction score*. Similarly, since Q(*P*) quantifies the error in terms of spatial coordinates, we call this quantity the *spatial coherence score*.

For the benchmark, we use the entropy regularized Wasserstein distance and thus solve a regularized linear Optimal Transport problem with *ε* =10^−1^ and report the linear term (without the entropy). This quantity is also called the Sinkhorn distance in the literature^77^. Benchmark plots were produced using the Seaborn package.

For the evaluation of the cell type transition accuracy, we used the gene expression predicted by each method at time *t*+ 1, then solved a regularized linear Optimal Transport problem between this predicted gene expression and the true measurements at *t*+ 1. The resulting transport plan was then used as a similarity matrix on which we applied 10-nearest neighbours classification. This resulted in cell type predictions which we compared with the ground truth. Importantly, those cell type predictions are only based on gene expression predictions and didn’t use spatial coordinates as an input, neither in the potential nor in the transport plan. The spatial information was only leveraged by STORIES during the training, and not at prediction time.

### Qualitative evaluation of trajectories

For Figures 2D-E and Extended Data Figures 2C-E & 3B-C & 4-6, we compared transport plans involved in STORIES and other methods which are used to match each method’s predictions with the true gene expression measurements at the next time point. Nonetheless these two experiments differ in an important way.

In Figure 2D-E, we show on one side the FGW transport plan used to compute STORIES’ training loss between the model’s predictions and gene expression measurements from the next time point. On the other side we show the Wasserstein transport plan used to compute the linear version of STORIES’ training loss.

In Extended Data Figures 2C-E & 3B-C & 4-6, we compute for both STORIES and PRESCIENT, the linear optimal transport plan between each model’s predictions and the gene expression measurements at the next time point. This allows us to compare the two methods only based on their output prediction without injecting spatial information at evaluation time.

Given a transport plan for each method, we compare how cells are matched at the cell type level. Formally, let us consider the indicator vector *a* ∈ ℝ^*n*^ where *a*_*i*_ = 1 if the *i* -th cell corresponds to a given cell type, and *a*_*i*_ = 0 otherwise. The transport plan ***P*** between the prediction *ρ*_*t*_ and the ground truth *μ*_*t*_ is applied to the indicator, yielding a vector b = *P*^⊤^ *a* representing the mass transported from a towards each cell in the second time point. These densities are plotted on top of the spatial coordinates for Figures 2D-E and Extended Data Figures 2C-E & 3B-C & 4-6.

### Computational growth rate

Growth rate estimation has only been performed in the two biological test cases (sections 3-4 of the results), where we analyzed a restricted number of cell types and could validate the biological coherence of the computed growth rate (see details in paragraphs below). The choice of not using growth rate estimation in the benchmark was also done to have a comparison with the state-of-the-art that is only focused on the quality of their underlying model. Indeed, other state-of-the-art methods sometimes use alternative ways to model the growth rate^14,16^ or do not consider it at all^83^. Furthermore, previous works tested gene sets to compute the growth rate in the case of mice and humans, but not of zebrafish and axolotl^13,16,21^. In addition, the growth rate estimation requires knowing a gene signature that is well established for some organisms, but might be missing for less studied ones. We thus kept uniform marginals for the benchmark in Section 2 of Results.

#### Weight of marginals

When comparing a prediction 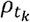 to the reference snapshot 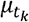, Yeo et al. proposes setting the weights *b*_*j*_ of the prediction 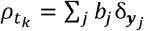 proportionally to a computationally derived growth rate. The motivation is that cells with a larger growth rate should be matched to more descendants. This idea was introduced in Waddington OT^13^ and recently reimplemented in MOSCOT^21^. In the next paragraphs, we follow MOSCOT’s implementation.

#### Proliferation and apoptosis

Computing the growth rate relies on cell-wise proliferation and apoptosis scores. We used Scanpy’s `sc.tl.score_genes` with lists of genes collected from the literature. The gene lists are described in more detail at the end of the section. We calculated gene scores on raw counts, after quality filtering of cells and genes.

#### Calculating the growth rate

For a given cell ***x***, let us call proli ∈ ℝ the proliferation score and apo_i_ ∈ ℝ the apoptosis score. We then define the birth rate *β*_*i*_ ∈ ℝ as

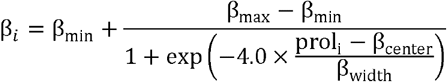

and the death rate γ_*i*_ ∈ ℝ as

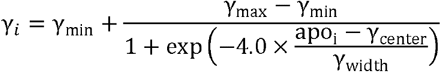

We used MOSCOT’s default parameters β_min_ γ_min_ 0.3, β_max_ γ_max_ 1.7, β_center_ 0.25, β_width_ 0.5, γ_center_ 0.1, γ_width_ 0.2. Finally, we defined the cell’s growth rate as

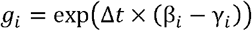

where Δ*t* is the time difference between populations. We obtained *a*_*i*_ by normalizing the growth rate

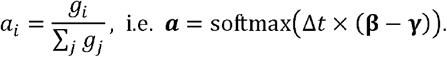

In this equation, *Δ*t plays the role of the softmax’s inverse temperature. The histogram *a* will thus be sharper for large values of Δ*t*. Most slices in our experiments were evenly sampled, so we set a fixed Δt = 1, which yielded sharp enough weight differences between cell types. In this estimation, Δ*t* plays the role of an inverse temperature parameter. A higher Δ*t* will result in a more entropic distribution of growth rates without changing the ranking of the cells in this distribution. On the other hand, using Δ*t* = 0 leads to estimating a constant growth rate for all cells which is therefore equivalent to not modeling growth. The growth rate models the potential division (rate > 1) or death (rate < 1) of cells between the measured time points. We relied on prior knowledge of the length of the cell cycle in the biological conditions we studied to make sure that our choice of Δ*t* was reasonable. In the axolotl regeneration case, for all tested values of Δ*t* our estimated growth rates were higher for less differentiated cells, which makes biological sense. In addition, we found in the literature that in the context of an injury, axolotl spinal cord cells might take between approximately 5 and 15 days to divide^84^. This means that cells could divide at most twice between consecutive timepoints in our dataset, thus giving birth to 4 descendants. As shown in Extended Data Figure 10A-C, using Δ*t* = 1 is coherent with this estimation as the maximum estimated growth rate is 4 in this case. While using a higher value for Δ*t* leads to unreasonably high estimated growth rates, using a value between 0 and 1 interpolates between not modeling growth and the upper bound on maximum growth between measured timepoints. In mouse midbrain development, we also observed that our estimated growth rates were higher for the less differentiated cells: radial glial cells. Moreover, we found in the literature that mouse cells between 13 and 17 embryonic days (in the same range as our dataset which has 3 time points at E12.5, E14.5 and E16.5) take between approximately 16 and 26 hours to divide^85^. This means that cells could replicate at most three times between consecutive timepoints in our dataset (which would result in a cell giving birth to 8 descendants). As shown in Extended Data Figure 10D, our estimated growth rates with Δ*t* = 1 are coherent since they are below the theoretical upper bound.

#### Dorsal midbrain

For Section 3 of Results, we used murine proliferation and apoptosis gene sets from MOSCOT. Proliferation genes come from https://doi.org/10.1038/nature20123 and apoptosis genes from https://www.gsea-msigdb.org/gsea/msigdb/cards/HALLMARK_P53_PATHWAY. See Supplementary Text 1 and Supplementary Text 2 for the gene names.

#### Axolotl neuron regeneration

For Section 4 of Results, we used an NSC axolotl gene set described in the original publication https://www.science.org/doi/10.1126/science.abp9444 to represent proliferation and a human apoptosis gene set from MOSCOT, originally from gsea-msigdb’s HALLMARK_APOPTOSIS. See Supplementary Text 3 and Supplementary Text 4 for the gene names.

### Trajectory inference

#### Gene imputation

In our analysis, the gene expression trends (Figure 3 and Figure 4) would be negatively affected by the sparsity of gene expression. Cellrank demonstrated good performances in identifying gene expression trends with MAGIC^26,86^. We thus applied MAGIC gene imputation after all other preprocessing steps. We computed the exponentiated Markov transition matrix on the Harmony-aligned PCA space instead of the original PCA. We did not use the imputed signal for tasks other than gene expression trends.

#### Potential visualization

The neural potential *J*_*θ*_ is a functional defined on the *d*-dimensional space of Harmony-aligned principal components. To visualize the potential as a Waddington-like landscape defined on 2 dimensions, we proceed similarly to Qin et al.^87^.

- First, we compute the potential *J*_*θ*_ (***x***) associated with each cell ***x***.
- Then, we use Scipy’s RBF interpolation and the 2-dimensional Isomap coordinates of the cells to define a potential on a 2-dimensional grid^88^.
- The cells are projected on the surface using the interpolator.
- Finally, the maximum value is thresholded.

We rendered the resulting surface and point cloud using Blender’s Python API.

#### Cell-cell transition matrix

We used CellRank’s VelocityKernel with a velocity *v*(*x*) =− ∇ *J*_*θ*_ (***x***) for a trained *J*_*θ*_. Based on this kernel, we computed a cell-cell transition matrix, and the trajectory plots Figure 3B, Figure 4B.

#### Cell fate probabilities

After computing the VelocityKernel, we use CellRank’s GPCCA estimator to compute cell fate probabilities. We set the number of macrostates to the number of cell types present in the dataset and set the terminal states to [“dpEX”, “nptxEX”, “mpEX”] for the axoltol and [“neuB”, “glioB”] for the mouse midbrain. Extended Data Figure 7A was obtained by aggregating those cell fate probabilities at the cell type level with the CellRank function *cellrank*.*pl*.*aggregate_fate_probabilities*.

#### Gene expression trends

We fit the MAGIC-imputed gene expression as a function of the learned potential, using Scipy’s Spline Regression^88^. To plot the gene expression cascades in Figure 3D, Figure 4D, we order genes by the value of potential for which the maximum regressed expression value is achieved. Genes are then split into equally sized groups illustrating different stages of differentiation (10 groups for Figure 3D and 2 groups for Figure 4D). Finally, regressed values for the genes with the best regression scores in each group are displayed (3 per group in Figure 3D and 15 per group in Figure 4D).

#### Transcription factor enrichment

We perform TF-target enrichment based on the TRRUST dataset, which contains “Activation”, “Repression”, and “Unknown” links based on a curated literature. For each TF in the database, we perform a one-sided Wilcoxon rank-sum test comparing the list of regression scores of its *n*_1_ target genes, and the list of regression scores of the other *n*_2_ genes. The number of target genes is different for each TF, but in all cases *n*_1_ + *n*_2_ =10,000. Figure 3H and Figure 4H display the TFs ranked by *p*-value.

## Data availability

1. We retrieved the mouse Stereo-seq atlas from Chen et al.^4^, available at https://db.cngb.org/stomics/mosta/.
2. We retrieved the zebrafish Stereo-seq atlas from Liu et al.^6^, available at https://db.cngb.org/stomics/zesta/.
3. We retrieved the axolotl Stereo-seq atlas from Wei et al.^7^, available at https://db.cngb.org/stomics/artista/.
4. We retrieved mouse proliferation genes from Tirosh et al.^89^ (see Supplementary Text 1) and apoptosis genes from gsea-msigdb’s HALLMARK_P53_PATHWAY (see Supplementary Text 2), available at https://www.gsea-msigdb.org/gsea/msigdb/human/geneset/HALLMARK_P53_PATHWAY.html.
5. We retrieved an NSC axolotl gene set from Wei et al.^7^ (see Supplementary Text 3) and a human apoptosis gene set from gsea-msigdb’s HALLMARK_APOPTOSIS (see Supplementary Text 4), available at https://www.gsea-msigdb.org/gsea/msigdb/human/geneset/HALLMARK_APOPTOSIS.html.

## Code availability

The Python package for STORIES is hosted at https://github.com/cantinilab/stories^90^. It can be installed easily by running “pip install stories-jax”. Code to reproduce the experiments and figures is available at https://github.com/cantinilab/stories_reproducibility/.

