## Supplementary material for "STORIES: learning cell fate landscapes from spatial transcriptomics": Supelementary text

### Supplementary Text 1: Mouse proliferation gene set

The following list of genes was retrieved from https://moscot.readthedocs.io under `utils.data.proliferation_markers`. The original source for these genes is [Tirosh et al., 2016].

| | | | | | | |
|---|---|---|---|---|---|---|
| Mcm4 | Ccnb2 | Gmnn | Wdr76 | Gins2 | Cdc45 | Nuf2 |
| Smc4 | Cenpa | Cdc6 | Ung | Cdc20 | Ckap5 | Anln |
| Gtse1 | Cenpe | Pold3 | Hn1 | Tpx2 | Ctcf | Nasp |
| Ttk | Cdca8 | Ckap2l | Cks2 | Rrm1 | Clspn | Hmmr |
| Rangap1 | Ckap2 | Fam64a | Kif20b | Cdc25c | Chaf1b | Rfc2 |
| Top2a | Rad51 | Ubr7 | Cdk1 | Cdca3 | Tacc3 | Prim1 |
| Hmgb2 | Pcna | Fen1 | Slbp | Ect2 | Mcm5 | Uhrf1 |
| Ccne2 | Ube2c | Bub1 | Aurkb | Pola1 | Anp32e | |
| G2e3 | Lbr | Brip1 | Kif11 | Ndc80 | Casp8ap2 | |
| Tmpo | Cenpf | Atad2 | Cks1b | Rpa2 | Tubb4b | |
| Nusap1 | Birc5 | Psrc1 | Blm | Nek2 | Kif23 | |
| Ncapd2 | Dtl | Rrm2 | Msh2 | Mki67 | Exo1 | |
| Mcm2 | Dscc1 | Tipin | Gas2l3 | Mcm6 | Tyms | |
| Kif2c | Cbx5 | E2f8 | Rad51ap1 | Cdca7 | Hjurp | |
| Aurka | Mlf1ip | Cdca2 | Usp1 | Dlgap5 | Hells | |

### Supplementary Text 2: Mouse apoptosis markers

The following list of genes was retrieved from https://moscot.readthedocs.io under `utils.data.apoptosis_markers`. The original source for these genes is the `HALLMARK_P53_PATHWAY` pathway from https://www.gsea-msigdb.org/.

| | | | | | | |
|---|---|---|---|---|---|---|
| Ercc5 | Serpinb5 | Krt17 | Pmm1 | Sphk1 | Cdkn2a | Nhlh2 |
| Pcna | Bmp2 | Inhbb | Steap3 | Dgka | Tpd52l1 | Def6 |
| Trib3 | Procr | Btg2 | Phlda3 | Cdkn2aip | Sesn1 | Cdkn2b |
| Blcap | Ada | Tnni1 | Rgs16 | Hmox1 | Foxo3 | Jun |
| Fgf13 | Irak1 | Ier5 | Slc19a2 | Rrad | Ddit4 | Slc35d1 |
| Tspyl2 | Sat1 | Adck3 | Ephx1 | Cdh13 | Zfp365 | Plk3 |
| Zmat3 | Hspa4l | Ptpn14 | Atf3 | Vwa5a | Osgin1 | Rnf19b |
| Slc7a11 | Tm4sf1 | Notch1 | Rxra | Zbtb16 | Cgrrf1 | Fuca1 |
| Rap2b | Fbxw7 | Ralgds | Ak1 | Rps27l | Abhd4 | Wrap73 |
| S100a4 | S100a10 | Stom | Ddb2 | Mapkapk3 | Kif13b | Rchy1 |
| Triap1 | Prkab1 | Cd82 | Il1a | Traf4 | Hint1 | Ip6k2 |
| Lrmp | Tm7sf3 | Trafd1 | Pom121 | Tgfb1 | Sertad3 | Pdgfa |
| Cebpa | Klk8 | Vamp8 | Retsat | Bax | Ppp1r15a | Tprkb |
| Rpl18 | Aen | Mxd1 | Sec61a1 | Rrp8 | Ccp110 | Xpc |
| Prmt2 | Nupr1 | H2afj | Ldhb | Btg1 | Ctsd | Mknk2 |
| Mdm2 | Cd81 | Dram1 | Hras | Ddit3 | Perp | Apaf1 |
| Gls2 | Rps12 | Ikbkap | Txnip | Cdk5r1 | Gm2a | Tcn2 |
| Ppm1d | Hist3h2a | Lif | Tsc22d1 | Rad51c | Alox8 | Upp1 |
| Tob1 | Trp53 | Ccng1 | St14 | Hexim1 | Fdxr | Cyfip2 |
| Plk2 | Sdc1 | Tax1bp3 | Gnb2l1 | Ccnk | Slc3a2 | Gpx2 |
| Jag2 | Fas | Zfp36l1 | Rad9a | Ndrg1 | Itgb4 | Fos |
| Plxnb2 | Rhbdf2 | Vdr | Baiap2 | Csrnp2 | Dcxr | Acvr1b |
| Sp1 | Ninj1 | Abat | Nol8 | Socs1 | F2r | Abcc5 |
| Cdkn1a | Trp63 | Tap1 | Fam162a | Ier3 | App | Polh |
| Dnttip2 | Clca2 | Wwp1 | Klf4 | Ankra2 | Ei24 | Hbegf |
| Sfn | Epha2 | Mxd4 | Iscu | Ccnd3 | Hdac3 | |

| | | | | | |
|---|---|---|---|---|---|
| Rb1 | Gadd45a | Tgfa | Ccnd2 | Rab40c | Ctsf |
| Ptpre | Eps8l2 | Nudt15 | Casp1 | Bak1 | Hist1h1c |

### Supplementary Text 3: Axolotl neural stem cell gene set

The following list of genes was retrieved from [Wei et al., 2022].

| | | | | |
|---|---|---|---|---|
| AMEX60DD001640 | AMEX60DD002950 | AMEX60DD005040 | AMEX60DD009861 | AMEX60DD011133 |
| AMEX60DD016851 | AMEX60DD022108 | AMEX60DD029807 | AMEX60DD031236 | AMEX60DD033253 |
| AMEX60DD034100 | AMEX60DD036933 | AMEX60DD038483 | AMEX60DD039579 | AMEX60DD041912 |
| AMEX60DD043225 | AMEX60DD043789 | AMEX60DD044011 | AMEX60DD044899 | AMEX60DD045542 |
| AMEX60DD045792 | AMEX60DD049907 | AMEX60DD050822 | AMEX60DD051580 | AMEX60DD052006 |
| AMEX60DD055073 | AMEX60DDU001006420 | | | |

### Supplementary Text 4: Human apoptosis gene set

The following list of genes was retrieved from https://moscot.readthedocs.io under `utils.data.apoptosis_markers`. The original source for these genes is the `HALLMARK_APOPTOSIS` pathway from https://www.gsea-msigdb.org/.

| | | | | | | | | | |
|---|---|---|---|---|---|---|---|---|---|
| ADD1 | AIFM3 | ANKH | ANXA1 | APP | ATF3 | AVPR1A | BAX | CASP2 | TNFSF10 |
| BCAP31 | BCL10 | BCL2L1 | BCL2L10 | BCL2L11 | BCL2L2 | BGN | BID | CASP3 | TOP2A |
| BIK | BIRC3 | BMF | BMP2 | BNIP3L | BRCA1 | BTG2 | BTG3 | CASP1 | TSPO |
| CASP4 | CASP6 | CASP7 | CASP8 | CASP9 | CAV1 | CCNA1 | CCND1 | CCND2 | TXNIP |
| CD14 | CD2 | CD38 | CD44 | CD69 | CDC25B | CDK2 | CDKN1A | SOD2 | VDAC2 |
| CDKN1B | CFLAR | CLU | CREBBP | CTH | CTNNB1 | CYLD | DAP | DAP3 | WEE1 |
| DCN | DDIT3 | DFFA | DIABLO | DNAJA1 | DNAJC3 | DNM1L | DPYD | EBP | XIAP |
| EGR3 | EMP1 | ENO2 | ERBB2 | ERBB3 | EREG | ETF1 | F2 | F2R | FAS |
| FASLG | FDXR | FEZ1 | GADD45A | GADD45B | GCH1 | GNA15 | GPX1 | GPX3 | |
| GPX4 | GSN | GSR | GSTM1 | GUCY2D | H1-0 | HGF | KRT18 | LEF1 | |
| HMGB2 | HMOX1 | HSPB1 | IER3 | IFITM3 | IFNB1 | IFNGR1 | LGALS3 | LMNA | |
| IGF2R | IGFBP6 | IL18 | IL1A | IL1B | IL6 | IRF1 | ISG20 | JUN | |
| LUM | MADD | MCL1 | MGMT | MMP2 | NEDD9 | NEFH | PAK1 | PDCD4 | |
| PDGFRB | PEA15 | PLAT | PLCB2 | PLPPR4 | PMAIP1 | PPP2R5B | RETSAT | RHOB | |
| RHOT2 | RNASEL | ROCK1 | SAT1 | SATB1 | SC5D | SLC20A1 | SMAD7 | SOD1 | |
| PPP3R1 | PPT1 | PRF1 | PSEN1 | PSEN2 | PTK2 | RARA | RELA | SPTAN1 | |
| SQSTM1 | TAP1 | TGFB2 | TGFBR3 | TIMP1 | TIMP2 | TIMP3 | TNF | TNFRSF12A | |
